# Dopaminergic tone predicts rhythmic temporal attention in Parkinson’s disease

**DOI:** 10.64898/2026.09.08.749528

**Authors:** Bar Yosef, Barathi Balamurugan, Mai Miura, Katy Cross

## Abstract

Beat perception, the ability to perceive and anticipate a regular pulse in music, is thought to depend on the basal ganglia. Dopaminergic signaling is a candidate mechanism given its prominence in basal ganglia circuitry and established function in encoding temporal predictions. Parkinson’s disease (PD) provides a model for studying dopaminergic contributions to beat perception. To test the hypothesis that higher dopaminergic tone is associated with better beat perception, we conducted a study in PD patients (n=55) and matched controls (n=53). Participants performed a musical beat perception task, the Beat Drop Alignment Test (BDAT). They also completed a non-timing control task measuring musical pitch perception, the Mistuning Perception Test (MPT). We identified a double dissociation in PD patients: daily dopamine dose was associated with beat perception (r=.37, p=.005) but not pitch perception (r=.12, p=.397), while musical background was associated with pitch perception (r=.55, p<.001) but not beat perception (r=.16, p=.236). Results remained consistent when controlling for disease duration and age. This dissociation suggests the effect of dopamine is specific to beat perception and not reflective of a general improvement in auditory processing. High-dopamine patients outperformed both low-dopamine patients and age-matched controls, and this difference was driven by better detection of on-beat than off-beat events. Together, these results suggest that dopamine is associated with enhanced beat perception, potentially through modulation of temporal attention to expected beat times.

## 1 Introduction

Stimuli in the real world often unfold rhythmically. Our brains take advantage of these temporal patterns to anticipate upcoming events. Executing motor actions, understanding language, and listening to music all involve the perception of temporal patterns. When we tap our feet or bob our heads to music, we are engaging in beat perception—the extraction and maintenance of a regular pulse from a musical rhythm. Beat perception is a predictive process since a regular beat can be perceived even when some beats are not marked by sound and some sounds occur off-beat. This capacity to anticipate and synchronize with a beat supports joint rhythmic behaviors such as chanting, drum circles, and dancing that have been observed in every human culture and are thought to promote social cohesion (Merker et al., 2009). Yet the neural mechanisms that give rise to this predictive ability remain incompletely understood.

Beat perception is thought to reflect strong and temporally precise connections between auditory and motor regions, with a number of studies showing activation of motor brain areas during rhythmic auditory processing (J. J. Cannon & Patel, 2021; Grahn & Brett, 2007; Zatorre et al., 2007). The basal ganglia in particular appear to play a crucial function in beat perception (Kasdan et al., 2022). They are involved in internal maintenance of the beat, as opposed to detection or tracking of an ongoing beat, which suggests a role in beat prediction (Grahn & Rowe, 2013). Dopamine, a key modulator of basal ganglia circuitry, is a potential substrate for instantiating these temporal predictions (Gershman & Uchida, 2019). Midbrain dopamine neurons encode not only the value, but also the timing of expected events (Schultz et al., 1997). Specifically, dopamine neurons’ response profile reflects how precisely the timing of an expected reward can be predicted (Fiorillo et al., 2008). In addition, dopamine depletion is associated with high levels of uncertainty about temporal predictions (Tomassini et al., 2016, 2019).

Dopamine has been proposed to encode precision as a gain that weights prediction errors, a mechanism shared with the attentional modulation of perception (Feldman & Friston, 2010; Friston et al., 2012); in this view, sharpening a temporal expectation and directing attention to it may reflect a common process. This literature, however, focuses on the preparation for single, discrete intervals rather than sustained periodic anticipation. Cannon (2021) suggested dopamine’s proposed precision-encoding role may extend to the periodic anticipation relevant for the recurring structure of a musical beat. His work builds on dynamic attending theory, which posits that attention entrains to the periodicity of external stimuli, forming “attentional energy pulses” that enhance perception at expected moments (Jones & Boltz, 1989; Large & Jones, 1999). Dynamic attending is reframed as continuous Bayesian inference in which dopamine signals the precision of temporal expectations. This computational framework proposes that the brain entrains to complex rhythmic auditory stimuli like music by producing ongoing estimates of the stimuli’s phase and tempo as they unfold. These timing expectations are compared to actual auditory input, and dopamine signals how much weight should be placed on the difference between the expectation and input, or error (J. Cannon, 2021). Putting this together, this attentional enhancement at expected beat times might be instantiated by dopaminergic signaling (J. Cannon, 2021; Friston et al., 2012; Large & Jones, 1999).

Experimental evidence for dopamine’s proposed contribution to rhythmic, as opposed to single-interval, temporal attention is lacking. To probe the acute effects of dopamine, researchers have compared Parkinson’s disease (PD) patients’ beat perception performance on and off dopaminergic medication. These studies have reported weak and inconsistent effects, including a marginal interaction in rhythm discrimination and no effect on a perceptual beat-alignment task (Cameron et al., 2016). Graded dopaminergic measures, which account for patients’ individual dopamine dose, have only been applied to interval timing or sensorimotor synchronization rather than perceptual beat perception . Beat perception studies in PD patients are critical to advancing understanding of dopaminergic contributions to rhythmic processing because sensorimotor synchronization studies are confounded by motor impairments in this population. To our knowledge, no study has related a continuous index of dopaminergic medication to beat perception ability.

In this study, we examined the relationship between dopamine intake and beat perception ability in PD patients and age-, sex-, and musical background-matched healthy controls (HC). Participants performed the Beat Drop Alignment Test (BDAT; Cinelyte et al., 2022), a musical beat perception task that requires internal maintenance of the beat to determine if a probe is on or off the beat. We hypothesized that participants with higher levodopa equivalent daily dose (LEDD; Jost et al., 2023) would be better at beat perception. To rule out LEDD improving music cognition broadly, we included a pitch perception control task (Larrouy-Maestri et al., 2019). Given dopamine’s proposed role in boosting temporal attention at expected beat times (J. Cannon, 2021; Friston et al., 2012; Large & Jones, 1999), we predicted that the association between dopaminergic medication and beat perception would be driven by on-beat trials specifically.

## 2 Methods

### 2.1 Participants

The study was conducted online and approved by the UCLA Institutional Review Board (IRB#24-000707); all participants provided informed consent. PD patients were recruited via email blasts through the UCLA Health patient portal (Epic, Verona, WI), flyers at the UCLA Movement Disorders clinic, and local Parkinson’s support group mailing lists. HC were recruited from flyers and mailing lists of older adults in the UCLA community and LA area.

Inclusion criteria were normal or corrected-to-normal hearing, no current hallucinations, age over 50, and no co-occurring neurological diagnoses. PD participants additionally required a PD diagnosis by a movement disorder specialist and no deep brain stimulation device, since the perturbation with stimulation may itself impact beat perception. Six PD and two HC participants were excluded due to low reported attention (i.e., responding “Some,” “Very little,” or “Almost no” to the question “How much attention did you give to the study?”). One PD was excluded for reporting that they were hard of hearing in a post-task debrief questionnaire. One HC was excluded for reporting confusion with the beat perception task instructions. One HC was excluded for reporting movement throughout the task. One PD was excluded for taking their dopaminergic medication more than 12 hours before testing. One PD patient was excluded for a presumed medication reporting error (a COMT inhibitor without a levodopa-containing medication).

The final sample included 55 PDs and 53 HCs matched in age, sex, and other demographic variables (Table 1). The PD group averaged an LEDD of 561.37 mg/day with a SD of 313.52.

**Table 1.**
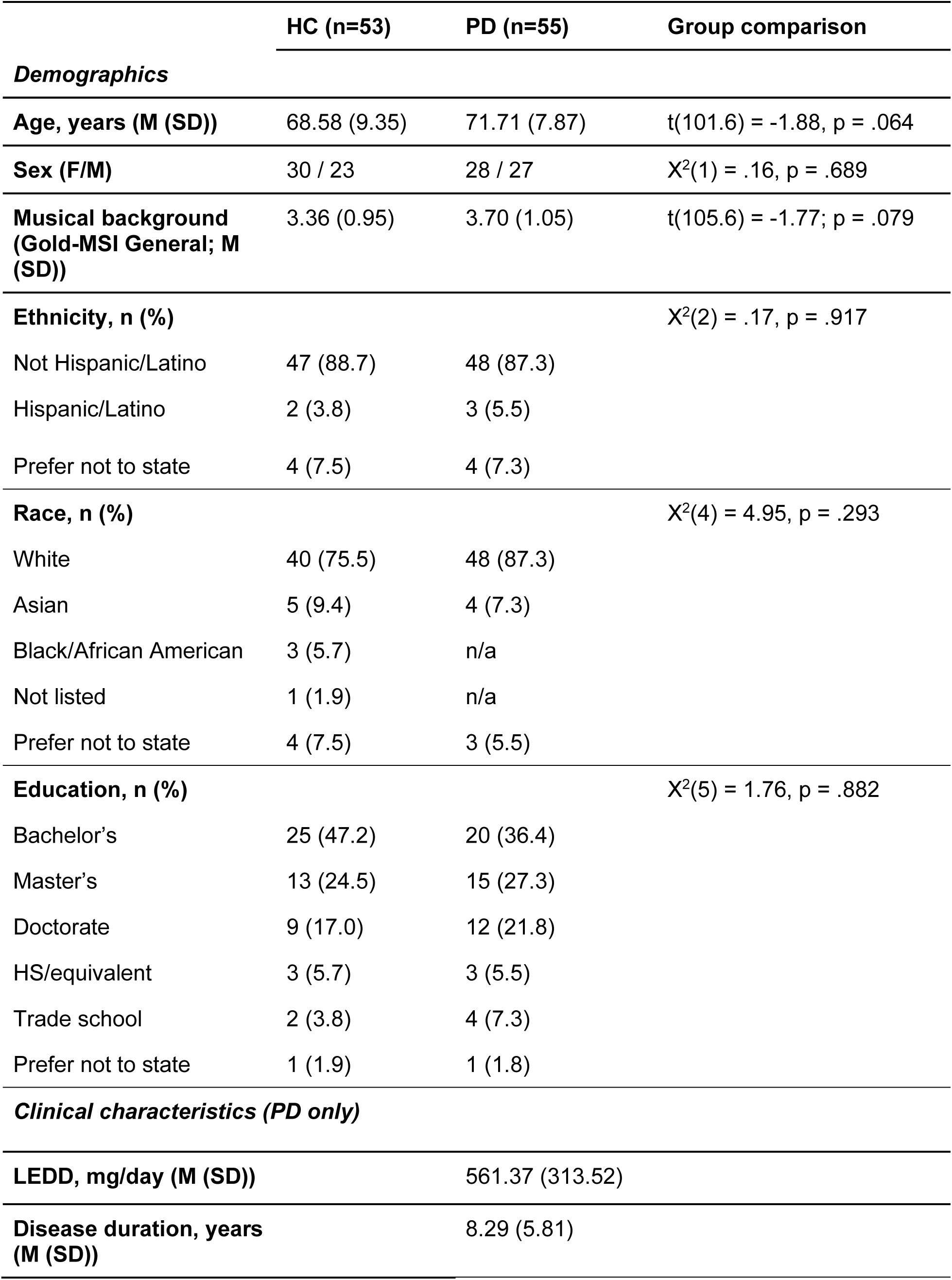
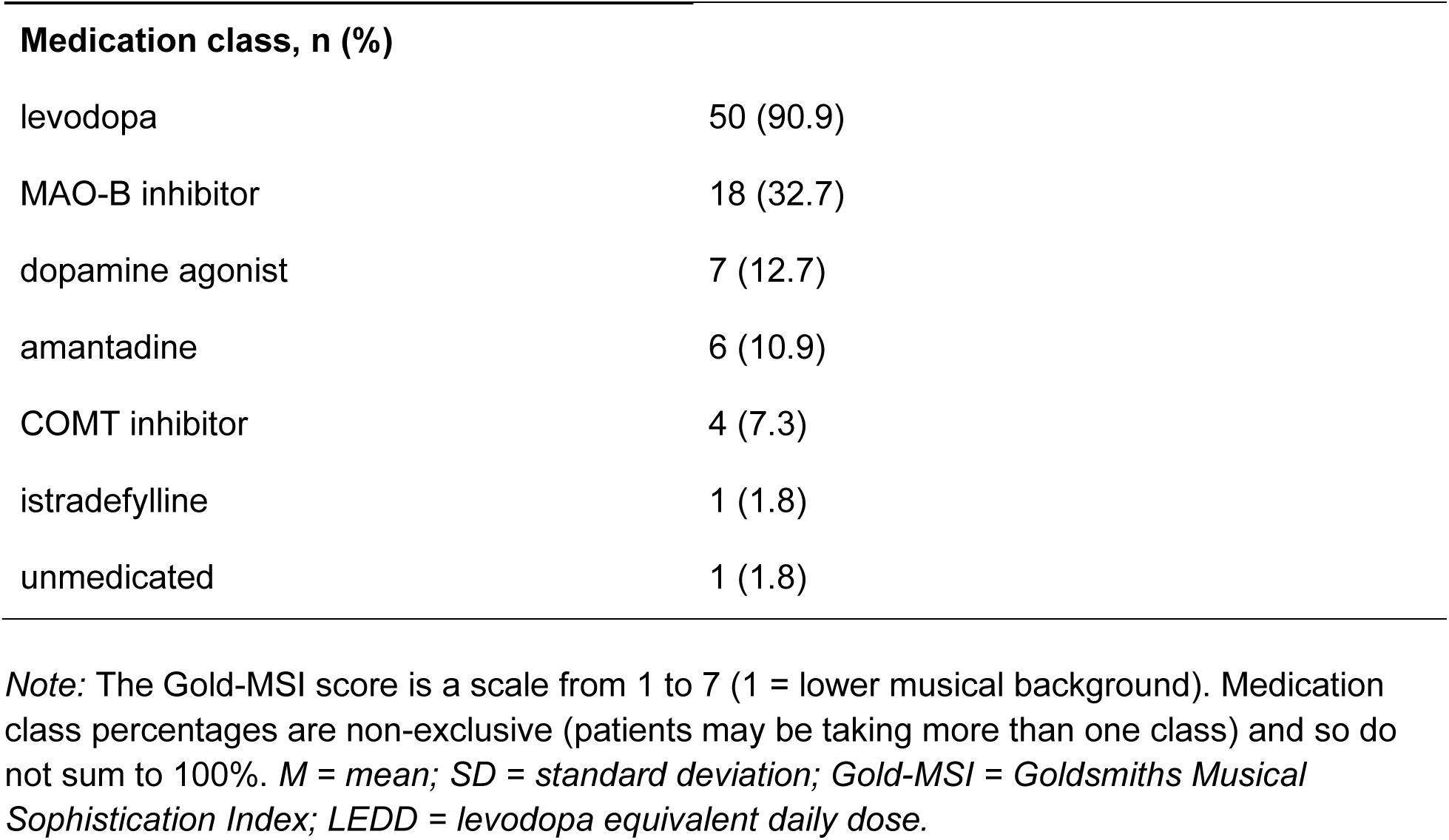
Participant demographics.

|  | HC (n=53) | PD (n=55) | Group comparison |
| --- | --- | --- | --- |
| **Demographics** |  |  |  |
| **Age, years (M (SD))** | 68.58 (9.35) | 71.71 (7.87) | $t(101.6) = -1.88, p = .064$ |
| **Sex (F/M)** | 30 / 23 | 28 / 27 | $X^2(1) = .16, p = .689$ |
| **Musical background (Gold-MSI General; M (SD))** | 3.36 (0.95) | 3.70 (1.05) | $t(105.6) = -1.77; p = .079$ |
| **Ethnicity, n (%)** | | | $X^2(2) = .17, p = .917$ |
| Not Hispanic/Latino | 47 (88.7) | 48 (87.3) |  |
| Hispanic/Latino | 2 (3.8) | 3 (5.5) |  |
| Prefer not to state | 4 (7.5) | 4 (7.3) |  |
| **Race, n (%)** | | | $X^2(4) = 4.95, p = .293$ |
| White | 40 (75.5) | 48 (87.3) |  |
| Asian | 5 (9.4) | 4 (7.3) |  |
| Black/African American | 3 (5.7) | n/a |  |
| Not listed | 1 (1.9) | n/a |  |
| Prefer not to state | 4 (7.5) | 3 (5.5) |  |
| **Education, n (%)** | | | $X^2(5) = 1.76, p = .882$ |
| Bachelor's | 25 (47.2) | 20 (36.4) |  |
| Master's | 13 (24.5) | 15 (27.3) |  |
| Doctorate | 9 (17.0) | 12 (21.8) |  |
| HS/equivalent | 3 (5.7) | 3 (5.5) |  |
| Trade school | 2 (3.8) | 4 (7.3) |  |
| Prefer not to state | 1 (1.9) | 1 (1.8) |  |
| **Clinical characteristics (PD only)** |  |  |  |
| **LEDD, mg/day (M (SD))** |  | 561.37 (313.52) |  |
| **Disease duration, years (M (SD))** |  | 8.29 (5.81) |  |

| **Medication class, n (%)** |  |
| --- | --- |
| levodopa | 50 (90.9) |
| MAO-B inhibitor | 18 (32.7) |
| dopamine agonist | 7 (12.7) |
| amantadine | 6 (10.9) |
| COMT inhibitor | 4 (7.3) |
| istradefylline | 1 (1.8) |
| unmedicated | 1 (1.8) |

*Note:* The Gold-MSI score is a scale from 1 to 7 (1 = lower musical background). Medication class percentages are non-exclusive (patients may be taking more than one class) and so do not sum to 100%. *M* = mean; *SD* = standard deviation; *Gold-MSI* = Goldsmiths Musical Sophistication Index; *LEDD* = levodopa equivalent daily dose.

### 2.2 Study Design

Participants performed an online task battery that included: the BDAT (Cinelyte et al., 2022), mistuning perception task (MPT, Larrouy-Maestri et al., 2019), and Goldsmiths Musical Sophistication Index (Gold-MSI) questionnaire (Müllensiefen et al., 2014).

The BDAT is a musical beat perception task. Participants were presented with a musical excerpt (stimulus presentation) and asked to track the beat. Following a brief pause in rhythmic events in the music (beat drop), they were asked to determine whether a single probe sound presented during the beat drop was on or off the beat (Figure 1A). The probe was presented either on Beat 3 or 4 or early or late relative to those beats (off beat). Off-beat probes were presented at seven displacement levels designed to have a linearly changing difficulty (15%, 18%, 20%, 23%, 26%, 31%, and 45% of the beat period; Cinelyte et al., 2022). Musical excerpts were approximately 11.5 seconds in duration and presented at five tempi ranging from 119 to 132 BPM (inter-beat intervals: 455-504 ms).

**Figure 1.**
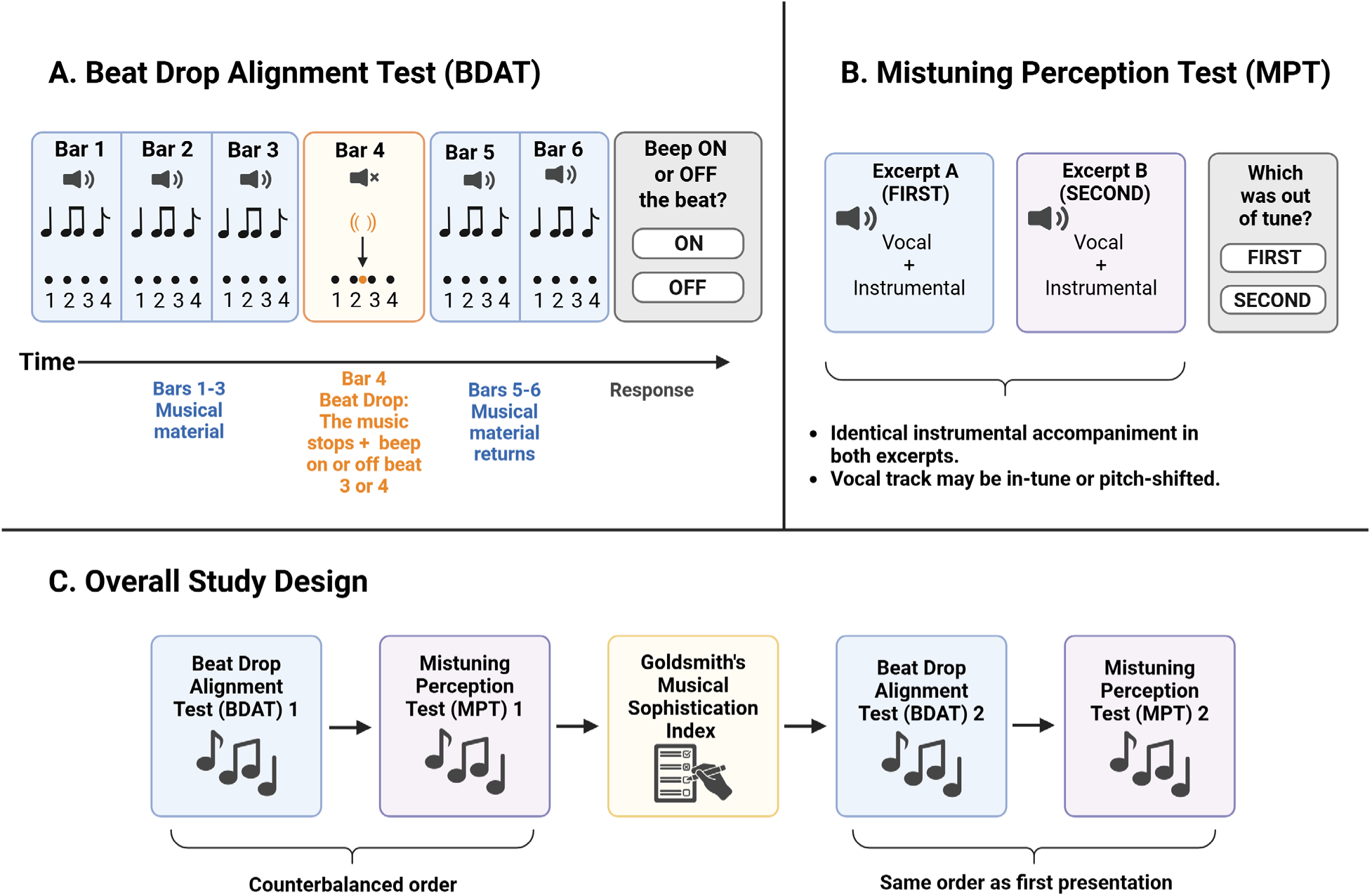
Structure of the Beat Drop Alignment Test (BDAT), Mistuning Perception Test (MPT), and overall study design. (A) In the BDAT, musical excerpts containing a beat (numbered dots in figure) are presented in bars 1-3 (box 1), followed by a beat-drop in bar 4 (box 2) when the music subsides. A probe (woodblock sound) is presented during the beat-drop bar on beat 3 or 4 (ON beat trials), or early or late relative to either beat (OFF beat trials, shown off beat 3 for illustration). The music returns in bars 5-6 (box 3), after which participants indicate whether the probe was ON or OFF the beat (box 4). (B) In the MPT, participants compare two excerpts with identical instrumental accompaniments (box 1-2) and determine whether the FIRST or SECOND excerpt contains the mistuned vocal track (box 3). (C) Participants completed the BDAT, MPT, and the Goldsmiths Musical Sophistication Index (Gold-MSI) followed by second repetitions of the BDAT and MPT.

The MPT is a musical pitch perception task (Larrouy-Maestri et al., 2019). Participants were presented with two musical tracks containing the same instrumental accompaniment. In one track, the vocalist is pitch shifted to be out of tune. Participants were asked to respond whether the vocalist is out of tune in the first or second track (Figure 1B). The musical excerpts ranged from 4.91 to 14.40 seconds in duration. The mistuned vocal track was shifted by 10-100 cents in 5-cent increments, being either sharp or flat relative to the instrumental accompaniment. The MPT was selected as a negative control (not expected to be related to dopamine) that has similar demands to the BDAT, including sustained attention to musical stimuli, auditory working memory, and decision-making with a binary response. Both tasks use item response theory (IRT)-based adaptive testing, in which item difficulty was selected according to participants’ estimated ability after each response (See Cinelyte et al. (2022) and Larrouy-Maestri et al. (2019) for more details). Adaptive testing allows for estimating a wide range of abilities in a short amount of time, and IRT is more flexible than simpler adaptive procedures like staircasing (Harrison & Müllensiefen, 2018; Magis & Barrada, 2017; Magis & Raîche, 2012).

The Gold-MSI is a 38-question survey that uses factor analysis to quantify musical background across five subscales: active engagement, musical training, emotions, singing abilities, and perceptual abilities. We report the “General Musical Sophistication” measure, a factor comprising 18 items selected for loading most strongly onto this general, higher-order factor (Müllensiefen et al., 2014).

Both the BDAT and MPT were performed twice. Each repetition was 25 trials long, and the adaptive procedure took place independently for each administration, so that the performance during the first administration did not influence item selection during the second administration. The Gold-MSI was performed in between first and second repetitions of both tasks (Figure 1C). The order of the BDAT and MPT was counterbalanced across participants, and the same for the second repetition within participants. Participants were asked not to move to the beat during the tasks because dopamine is involved in movement, and we did not want this to impact task performance. The battery took about 1 hour.

We also included a post-task questionnaire that asked participants to report how much attention they gave to the study (on a scale from 1 to 5, with 1 being “almost none” and 5 being “a lot”) and how difficult they found the BDAT and MPT (on a scale from 1 to 7, with 1 being “very easy” and 7 being “very difficult”). The task battery was administered using shiny R on the German Society for Music Psychology (DGM) server (Chang et al., 2026). The eligibility, screening, and demographics questionnaires were administered on REDCap (Harris et al., 2009, 2019).

Patients who reported taking medications for Parkinson’s disease were presented with a list of common PD medications (both brand and generic names) and asked to check off all medications they were taking. For each checked medication, a list of the available doses specific to the medication was populated and patients checked which dose(s) they took followed by the number of pills taken per day. This structured assessment was designed to maximize accuracy of self-reported medication intake. We calculated LEDD according to conversion factors from Jost et al. (2023). LEDD is reported in units of 100 mg/day in regression results.

### 2.3 Analysis

Group differences in demographic and clinical characteristics (Table 1) were assessed using Welch’s t-tests for continuous variables (age, musical background) and chi-square tests of independence for categorical variables (sex, ethnicity, race, education).

Beat perception and pitch perception ability scores were derived using item response theory as in Cinelyte et al. (2022) and Larrouy-Maestri et al. (2019). Ability is analogous to a z-score, in which 0 represents the average ability of a normative sample and higher values indicate better performance. We averaged ability scores across both repetitions for each task, except for cases in which participants reported moving to the beat in the second repetition (n=3 HC and 1 PD). We used Pearson r correlation coefficients to examine the relationship of beat and pitch perception with LEDD. To ensure the robustness of our findings, we performed linear regressions adjusting for age and disease duration. We investigated a potential association between acute dopamine level and performance by correlating time since the most recent dose with task performance. Because time since the most recent dose was positively skewed, we used Spearman rank correlations as opposed to Pearson correlations. To compare low- and high-dopamine PD groups to healthy controls, we used a median split that separated patients into low (<500 mg, n=27) and high (≥500 mg, n=28) LEDD groups. We compared beat and pitch perception ability between groups using a one-way ANOVA and pairwise t-test comparisons. Self-reported attention and task difficulty were compared across the three groups (HC, low-LEDD PD, high-LEDD PD) using Kruskal-Wallis tests.

Finally, we performed a trial-specific analysis to determine whether the group differences in beat perception ability were driven by better performance for on-beat trials, which would suggest that dopamine boosts temporal attention at the expected beat time. We repeated the group comparisons separately for trials in which the probe was on the beat (“on-beat”) and trials with the probe off the beat (“off-beat”) using pairwise t-tests. For these analyses, ability was recalculated separately for on and off trials using the same approach as the overall ability scores (Cinelyte et al., 2022; Magis & Barrada, 2017).

All pairwise group comparisons were corrected for multiple comparisons using the Benjamini-Hochberg false discovery rate (FDR) procedure to avoid inflating Type I error across the family of comparisons for that measure (Benjamini & Hochberg, 1995). The significance level was set at .05. All analyses were conducted in Python (version 3.11.4), using pandas and numpy for data handling, scipy.stats for statistical tests, and statsmodels for linear regression and false discovery rate correction. Figures were generated using matplotlib and seaborn.

## 3 Results

### 3.1 Dopamine dose and beat perception ability in PD

First, we examined whether there was an association between LEDD and beat perception ability in PD patients. To ensure that LEDD does not simply result in improvement in general cognitive ability or attention, we also examined whether there was an association between LEDD and pitch perception ability. We identified a double dissociation: daily dopamine dose was associated with beat perception (r = .37, p = .005) but not pitch perception (r = .12, p = .397), while musical background was associated with pitch perception (r = .55, p < .001) but not beat perception (r = .16, p = .236; Figure 2). These associations remained significant in a full regression model controlling for age and disease duration (LEDD → beat: β = 0.125, p = .008; Gold-MSI → pitch: β = 0.348, p < .001), with neither predictor crossing over to the other outcome (LEDD → pitch: p = .430; Gold-MSI → beat: p = .204; Table 2).

**Figure 2.**
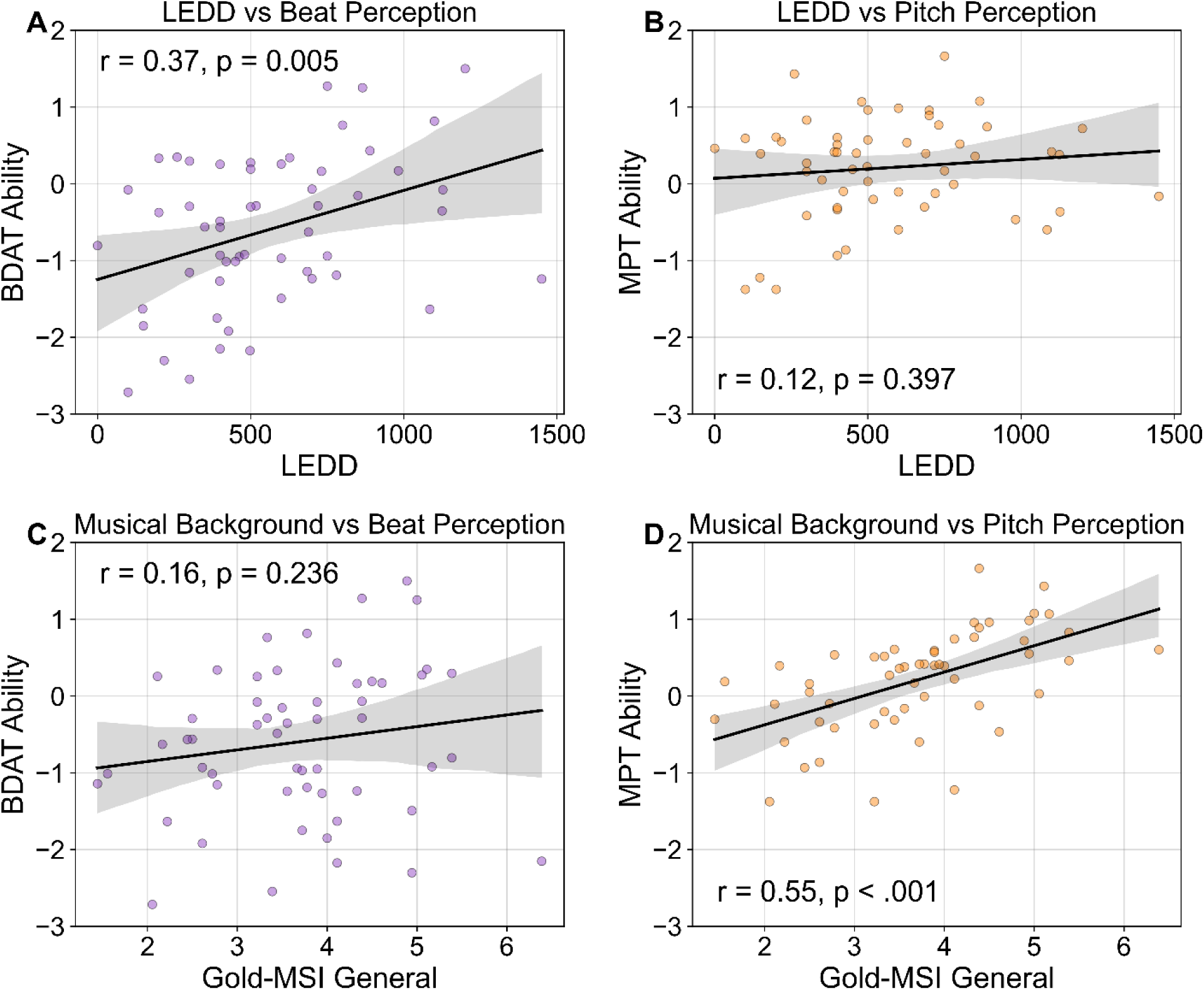
Dopamine dose has a specific association with beat perception in PD (n=55). (A) LEDD is associated with beat perception (BDAT) ability but (B) not pitch perception (MPT) ability, while (C) musical background (Gold-MSI General) is associated with pitch perception ability but (D) not beat perception ability, demonstrating a double dissociation. Line = best fit; shaded region = 95% CI.

**Table 2.**
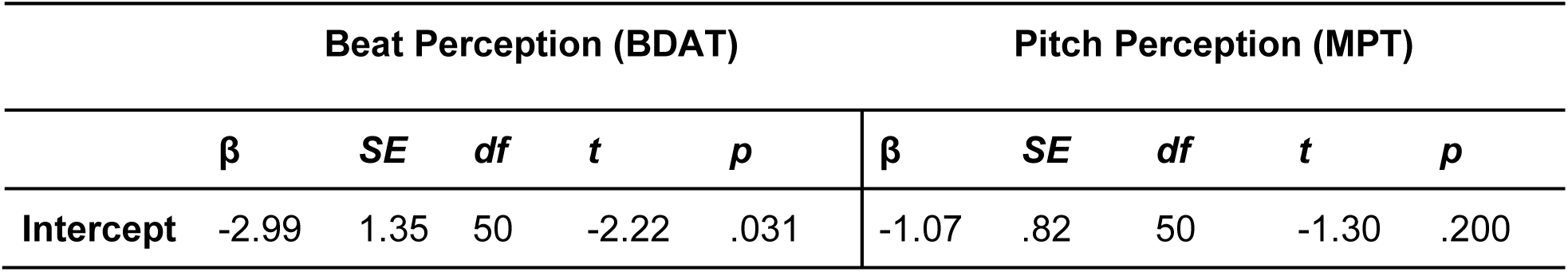
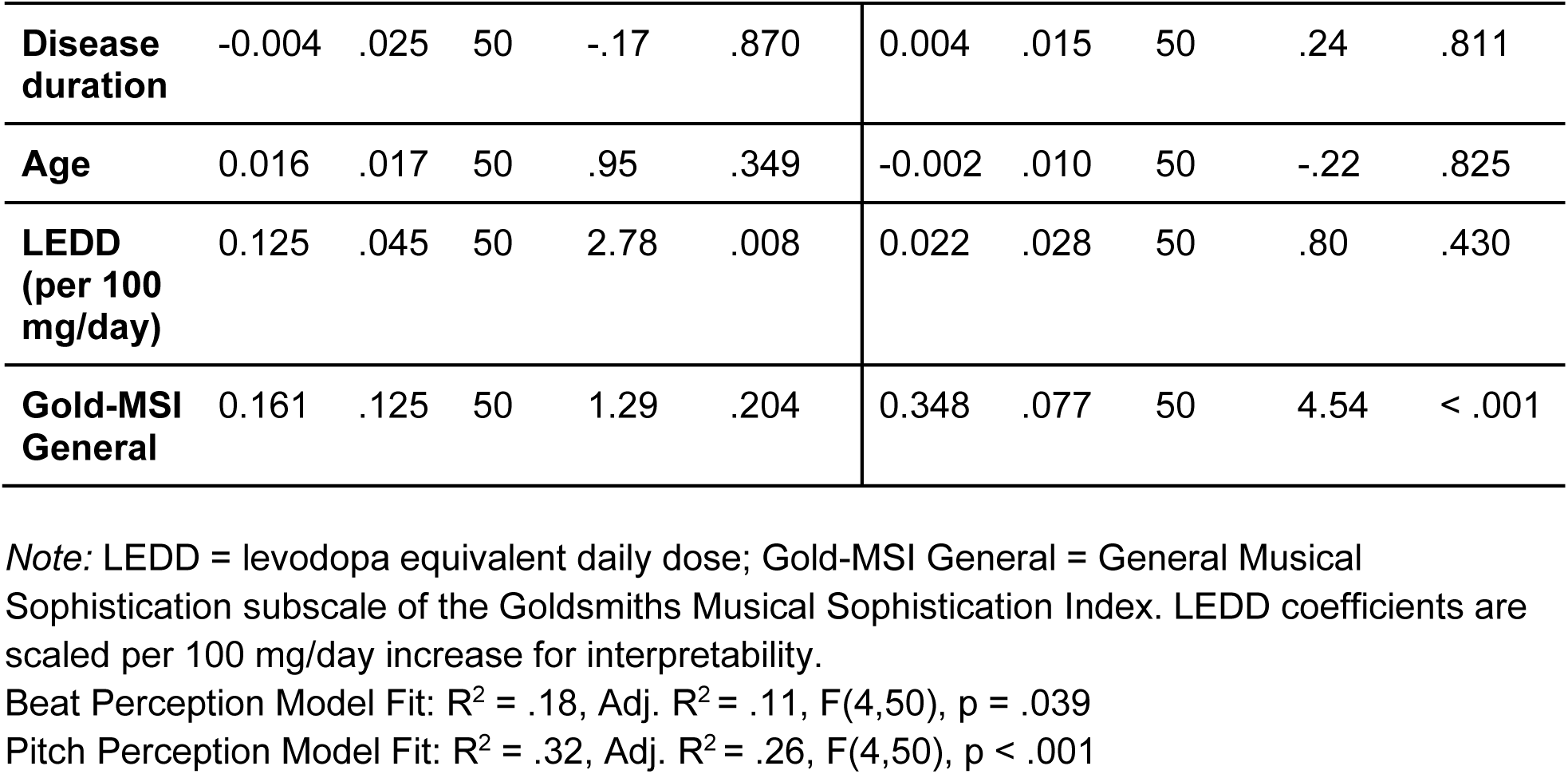
Multiple linear regression models predicting beat perception (BDAT) and pitch perception (MPT) ability in PDs from dopamine dose and musical background, controlling for age and disease duration (n=55).

A similar pattern between musical background and performance in the tasks was observed in healthy controls, with Gold-MSI having a significant correlation with MPT (r = .38, p = .0048) but not BDAT ability (r = .22, p = .11).

Disease duration was associated with LEDD (r = .4, p = .003; Supplemental Figure 1), consistent with disease severity. Importantly, disease duration was not associated with BDAT (p = .432) or MPT ability (p = .555) (Supplemental Figure 1), ruling out disease severity as a confound. In addition, we found that time since the patients’ most recent dose of Parkinson’s medication had no relationship with beat or pitch perception ability (BDAT: Spearman r = .03, p = .835; MPT: Spearman r = -0.14, p = .281).

### 3.2 PD patients on a high and low dopamine dose versus controls

Next, we examined group differences in BDAT and MPT ability across three groups: HC (n = 53), low-LEDD PD (n = 27), and high-LEDD PD (n = 28). For beat perception, we found a main effect of group in a one-way ANOVA (F(2,105)=6.75, p=.0017). High-LEDD PD patients (M = −0.16, SD = 0.86) outperformed both low-LEDD PD patients (M = −1.05, SD = 0.91; p = .002) and HC (M = −0.80, SD = 0.98; p = .006) on the BDAT, while low-LEDD PD and HC did not differ (p = .262; Figure 3A). No group differences were observed for pitch perception (no main effect of group in a one-way ANOVA, F(2,105)=2.27, p=.108). Group means were comparable across groups (HC: M = −0.03, SD = 0.78; low-LEDD: M = 0.08, SD = 0.73; high-LEDD: M = 0.33, SD = 0.58; Figure 3B).

**Figure 3.**
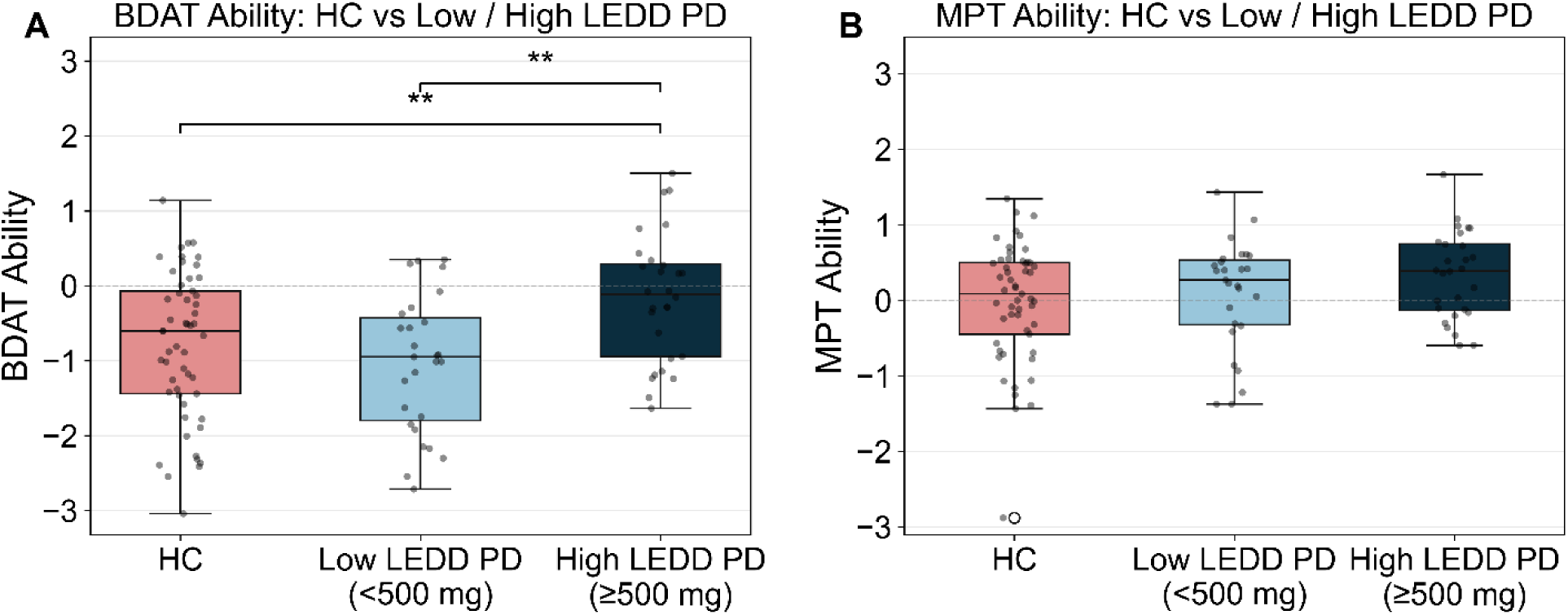
PD patients on a high dopamine dose outperform low-dose patients and age-matched healthy controls. (A) Beat perception ability (BDAT) and (B) pitch perception ability (MPT) across healthy controls (HC, n=53), low-LEDD PD (n=27), and high-LEDD PD (n=28). Boxplots show median and interquartile range; individual points represent subject-level scores. Significance bars reflect FDR-corrected pairwise comparisons within each measure (**p<.01).

Importantly, attention ratings did not differ across groups (HC: M = 4.51, SD = 0.50; low-LEDD PD: M = 4.56, SD = 0.51; high-LEDD PD: M = 4.46, SD = 0.51; H(2) = 0.45, p = .80). Self-reported task difficulty was also comparable across groups for both the BDAT (HC: M = 5.00, SD = 1.52; low-LEDD PD: M = 5.26, SD = 1.10; high-LEDD PD: M = 4.71, SD = 1.61; H(2) = 1.11, p = .574) and the MPT (HC: M = 4.98, SD = 1.26; low-LEDD PD: M = 5.30, SD = 1.27; high-LEDD PD: M = 4.93, SD = 1.25; H(2) = 2.46, p = .292).

### 3.3 Dopamine dose and on versus off beat trials

Finally, we analyzed trials in which the probe was on and off the beat separately in order to determine whether the performance differences were driven by on-beat trials, which would suggest that dopamine boosts temporal attention to the beat (J. Cannon, 2021; Friston et al., 2012). Between-group comparisons (Welch’s t-tests, FDR-corrected jointly across on- and off-beat conditions) showed that high-LEDD PD had higher ability than low-LEDD PD specifically for on-beat trials (p = .003). No other pairwise comparisons reached significance after correction; high-LEDD PD showed marginally higher on-beat performance than HC, significant before correction (p = .037) but not after FDR correction (p = .074; Figure 4). This pattern also held when on- and off-beat performance was quantified using raw accuracy (proportion correct) instead of ability, with high-LEDD PD again showing higher accuracy than low-LEDD PD specifically for on-beat trials (p = .026). These results suggest that the group difference in beat perception was primarily driven by on-beat trials.

**Figure 4.**
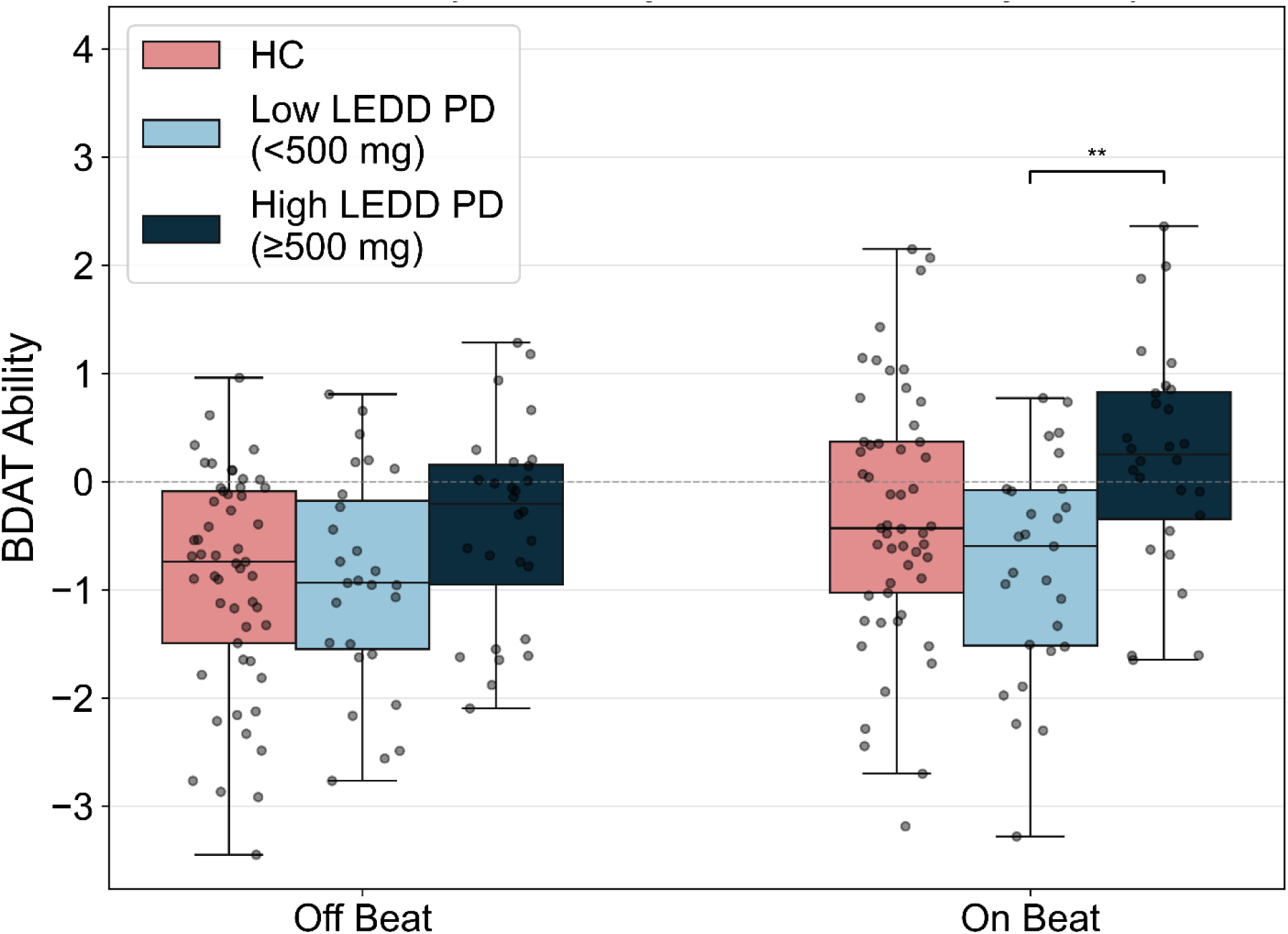
PD patients on a high dopamine dose have higher ability than low-dopamine dose patients in on-beat trials and not off-beat trials. Boxplots show median and interquartile range; individual points represent subject-level values. Significance bars reflect FDR-corrected pairwise comparisons between groups within each measure and between conditions within each group (**p<.01).

## 4 Discussion

The basal ganglia have been proposed to play an important role in beat perception (Kasdan et al., 2022). Beat perception is a form of relative timing and refers to the extraction and maintenance of a regular pulse from a musical rhythm. Basal ganglia involvement dissociates beat-based timing from other forms of interval timing (Teki et al., 2011) and is specifically associated with beat prediction as opposed to other cognitive processes (Grahn & Rowe, 2013). Given its role in encoding temporal predictions and prominence within basal ganglia circuitry, dopamine might underlie basal ganglia involvement in beat perception (Coull et al., 2011). In this study, we examined the relationship between total daily dopamine dose and beat perception performance in PD. We found that dopamine dose was associated with improved beat perception ability and had no relationship with pitch perception ability. This effect was driven by a higher performance for on-beat trials in high-dopamine PD patients, suggesting that dopamine is particularly important for temporal attention. Notably, high-dopamine PD patients outperformed both low-dopamine PD patients and age-matched controls. To our knowledge, this is the most specific evidence to date for a key function of dopamine in beat perception.

Although dopamine was only associated with beat perception and not pitch perception, we did find a relationship between musical background and pitch perception. This double dissociation suggests that the positive effect of dopamine is specific to rhythmic temporal processing. As musical background is related to pitch perception, the lack of correlation between dopamine dose and pitch perception is unlikely to be due to the invalidity of MPT. Musical background may have had a weak relationship to beat perception in our study because participants were asked not to move to the beat, which could make the strategies used to perform the task less related to how music is performed and enjoyed in the real world (Morillon & Baillet, 2017; Su & Pöppel, 2012). This pattern of results, with musical background being associated with pitch but not beat perception, was also found in our healthy control group.

Dopamine dose specifically predicted on-beat trial performance and not off-beat trial performance. This selectivity is informative about the underlying mechanism. One possibility is that dopamine reduces noise in the internal representation of elapsed time (Buhusi & Meck, 2005). However, such global sharpening would be expected to improve discrimination across the whole rhythmic sequence, rather than selectively at the beat. A more consistent account is that dopamine weights the precision of temporal predictions rather than the fidelity of the clock itself. Under Bayesian inference, dopamine encodes the precision of prediction errors, implemented as a gain on postsynaptic responses, and this precision-weighting is a mechanism shared with the attentional modulation of sensory processing (Friston et al., 2012). Because precision is allocated to specific predicted events rather than applied uniformly, its enhancement would manifest at moments where a temporal prediction exists, namely at the beat. This maps onto dynamic attending theory, under which attention is entrained to a periodic structure and heightened at expected times (Bouwer & Honing, 2015; J. Cannon, 2021; Large & Jones, 1999). Consistent with a role for dopamine in temporal attention, dopaminergic medication modulates performance of PD patients on the attentional blink task, which requires detecting two temporally-close targets (Slagter et al., 2016). Under a dynamic attending framework, attentional energy is entrained to the beat, heightening sensitivity at expected moments; a dopamine-driven enhancement of this process would manifest specifically at on-beat times, consistent with our results (Figure 5). To our knowledge, this link between dopaminergic precision and rhythmic temporal attention has not been directly tested, and our finding provides initial behavioral evidence for it.

**Figure 5.**
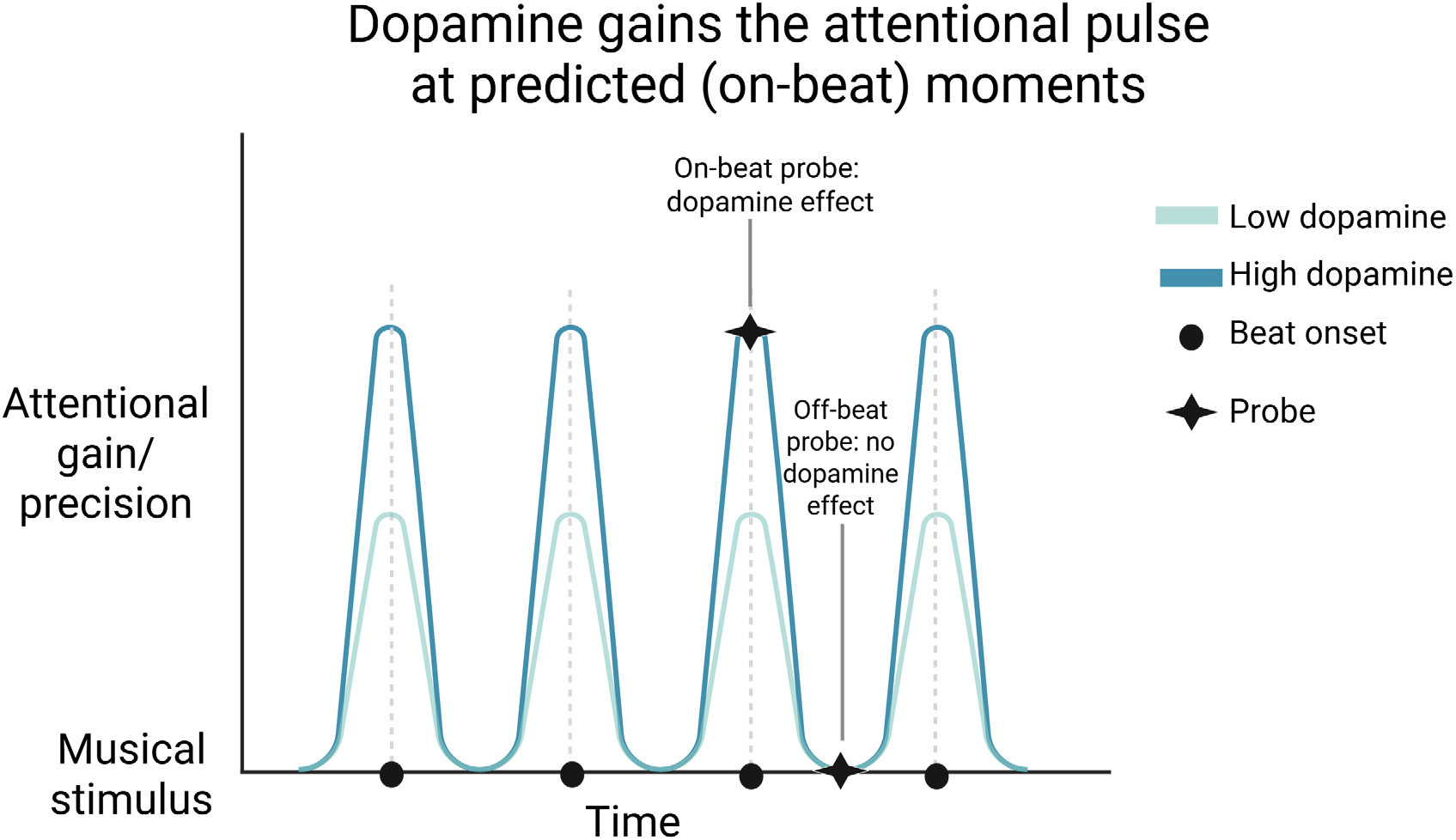
Schematic representation of the interpretation of our findings. Higher performance in the high-dopamine PD group was driven by on-beat trials. This might be due to dopamine boosting attentional gain at the beat onset (Friston et al., 2012; Large & Jones, 1999; Tomassini et al., 2016). Since this proposed mechanism boosts attention in a temporally precise manner specific to beat times, it is consistent with the lack of a difference between the high- and low-dopamine patients for off-beat trials.

Interestingly, high-dopamine PD patients outperformed healthy controls. This is consistent with evidence that PD patients have relatively intact dopamine levels in the ventral striatum, which receives projections from the ventral tegmental area, and decreased dopamine levels in the dorsal striatum, which receives projections from the substantia nigra pars compacta (Kish et al., 1988). Although PD patients are prescribed dopaminergic medications to offset loss of motor function driven by decreased dopamine in the dorsal striatum, the drugs also elevate dopamine in the relatively spared ventral striatum. Thus, PD patients can be “overdosed” in ventral striatum dopamine relative to healthy adults (Cools et al., 2002; Vaillancourt et al., 2013). Better beat perception in high-dopamine PDs compared to healthy older adults in our sample might be driven by medicated PDs having elevated dopaminergic processing in the ventral striatum supporting higher precision. This suggests that the ventral striatum may be involved in beat perception as opposed to the dorsal striatum and is consistent with recent experimental and theoretical accounts of ventral striatum involvement in temporal processing (J. Cannon, 2021; Takahashi et al., 2016). Since the ventral striatum is involved in reward processing, temporal processing and reward processing may rely on shared dopaminergic pathways (Braun Janzen & Thaut, 2019; Lake & Meck, 2013).

Research on time perception in PD as a model of basal ganglia dysfunction has a rich history. At the same time, there have been mixed findings on whether or not PD patients have impaired time perception (see Buikema et al. (2026) for a recent review). This picture is complicated by the fact that motor deficits in PD may confound studies that use motor responses, such as tapping along with an auditory tone and maintaining the rhythm after the tone stops (e.g., Singh et al., 2021). Beat perception studies are a valuable contribution to this literature because they enable studying temporal processing without a motor output. Yet, the several studies that have examined beat perception in PD have also yielded somewhat mixed results. For example, although Grahn & Brett (2009) found a deficit in discrimination of beat-based rhythms in PD, a follow-up study found no significant difference between PD patients and controls on a perceptual beat-alignment task similar to the one used in the current study (Cameron et al., 2016). Cameron et al. (2016) also compared PD patients on and off dopaminergic medication, although they did not find consistent results, with only a weak effect in a beat discrimination task and no effect on a beat-alignment task. Notably, this study did not report or account for the dosage of dopaminergic medication that patients were taking. Patients’ individual dopamine dose, which is known to have a complex relationship with a wide range of cognitive functions (Vaillancourt et al., 2013), might explain heterogeneity in beat perception in PD patients.

On the other hand, we may have found an effect of dopamine on beat perception in our study due to cumulative effects of dopaminergic medication that are not captured in acute on/off comparisons. This is supported by the null effect of time since medication on beat perception ability in this study. The tonic effect of dopamine might be related to the neurotransmitter’s role in plasticity at corticostriatal synapses (Calabresi et al., 1997; Sidhu et al., 2004). This is consistent with recent evidence that reinforcement learning mediated by basal ganglia dopamine activity drives learning of complex natural behaviors across development, such as vocal song learning in zebra finches (Kasdin et al., 2025). Yet another reason for the difference in our findings compared to Cameron et al. (2016) is that we used the BDAT, which requires internal maintenance of the beat during the beat drop, while their study used the beat alignment test (BAT), in which the probes are overlaid with the musical excerpt. Despite finding no effect of disease or dopaminergic medication on the BAT, their group has found a deficit in PD for a rhythm discrimination task that does require internal beat maintenance (Grahn & Brett, 2009). Taken with recent neuroimaging studies showing a specific role for the basal ganglia in internal maintenance of the beat (Grahn & Rowe, 2013; Nozaradan et al., 2017; Schwartze et al., 2011), effects of dopamine on beat perception may be better detected in tasks with greater temporal prediction demands.

Several limitations of this study should be acknowledged. Because LEDD is a correlational index of cumulative dopaminergic exposure rather than an experimental manipulation, we cannot establish that dopamine causally drives beat perception. LEDD is also associated with other disease variables, including severity, which we could not measure in this study due to it being conducted online. While the relationship between LEDD and beat perception ability remained after controlling for disease duration and age, LEDD remains an imperfect proxy for dopaminergic tone, and unmeasured factors such as medication responsiveness or patient symptoms could contribute. Notably, more advanced disease would, if anything, be expected to impair performance, making it unlikely that the positive association reflects general disease progression. This is supported by the lack of an association between disease duration, a proxy for disease severity, and beat perception ability. Together with the effect’s specificity to beat over pitch perception, this argues against a nonspecific explanation.

We also lacked a dopaminergic index in controls. Thus, the proposal that older healthy controls sit below a ventral striatum-driven beat perception optimum remains an interpretation permitted by knowledge of differential degeneration of the striatum in PD rather than one that our data establish (Kish et al., 1988). In addition, LEDD collapses drugs with differing receptor profiles and pharmacokinetics into a single scalar, which may obscure the mechanism.

Despite these constraints, our findings show that cumulative dopaminergic load tracks beat perception specifically. This relationship is carried by heightened ability for on-beat trials, and more heavily medicated patients can outperform matched controls. Together, these results position dopamine as a candidate mechanism linking basal ganglia function to the temporal allocation of attention in rhythm perception.

## CRediT authorship contribution statement

**Bar Yosef:** Conceptualization, Data curation, Formal analysis, Funding acquisition, Investigation, Project Administration, Methodology, Visualization, Writing – original draft, Writing – review and editing. **Barathi Balamurugan:** Data curation, Software, Formal analysis, Visualization. **Mai Miura:** Investigation, Project administration, Writing – review and editing. **Katy Cross:** Conceptualization, Funding acquisition, Methodology, Resources, Supervision, Writing – review and editing.

## Acknowledgements

We are grateful for the participants in this study, without whom this research would not be possible. We thank Dr. Klaus Frieler and the German Society for Music Psychology (DGM) for their support in setting up and hosting the online task battery. We would also like to thank Dr. Ramesh Balasubramaniam for guidance on our experimental design, Dr. Dean Buonomano for providing insights on analytic approaches and interpretations of our findings, Dr. Leo Ma for feedback on writing and interpretations, and Navya Garg for proofreading. This material is based upon work supported by the National Science Foundation Graduate Research Fellowship Program under Grant No. (DGE-2034835; DGE-2444110). Any opinions, findings, and conclusions or recommendations expressed in this material are those of the author(s) and do not necessarily reflect the views of the National Science Foundation. The graphical abstract and Figures 1 and 5 were created with BioRender.com.

## Declaration of generative AI and AI-assisted technologies in the manuscript preparation process

During the preparation of this work, the authors used Claude by Anthropic to program subsets of the analysis code in Python and to reorganize the data processing pipeline. This AI tool was also used to proofread the manuscript, such as to make sure values were reported correctly. After using this tool, the authors reviewed and edited the content as needed and take full responsibility for the content of the published article.

